# A Distinct Mode of Chromosome-End Genome Instability Revealed by Circulome Characterization and Analyses

**DOI:** 10.64898/2026.11.03.763977

**Authors:** Grant Roy, Zhixiang Lei, Matthew J. McCoy, Aysha Al-Sardar, Jerry Shay, Peter C. Scacheri, Yair Lotan, Stephen Levene, Andrew Fire, Massa J. Shoura

## Abstract

Genome instability is a central hallmark of cancer and it is characterized largely from sequence: mutation, copy-number change and rearrangement scored against a linear reference. The genome has additional degrees of freedom, such as the physical form (topology) of the DNA itself. A specific mode of localized genome dynamics changes both dimensions (sequence and topology) at once: defined chromosomal regions are also represented as extrachromosomal circular DNA (eccDNA). The biogenesis of eccDNA is a feature of many cancers, supplying the tumor with genetic elements that can participate in phenotypic selection, and supplying the observer with a window into the genomic instability landscape. Genome-wide maps of where these circles arise have not been available at scale or at high resolution. Building such maps requires physical selection of circular species, because most sequenced reads from unselected DNA derive from unrearranged sequence, conflating the signal from dynamic rearrangements with that from bulk chromosomal DNA. Here we combine biophysical and biochemical isolation of circular DNA with next-generation sequencing, applying it to a panel of cancer cell lines and to patient tumors, with non-cancerous cells and adjacent normal tissues as controls. The resulting genome-wide maps of the circular-element repertoire, the Circulome, reveal a consistent and previously unreported enrichment of these elements at particular chromosome ends. The circles map to regions extending well past the telomeres, reaching varying distances up to tens of megabases inward. This activity is accompanied by corroborating rearrangements within the linear genome, consistent with regions of chromosomal instability. Together these features define a spatially localized mode of instability in cancer, which we term Erosion at Distal Genomic Ends (EDGE). We formulate a classification problem to discriminate bladder tumor from matched normal tissue, finding that the EDGE signal outperforms the mean of all interior chromosomal regions, for each chromosome. These findings define a distinct spatial signature of cancer chromosome instability detectable through Circulome analysis, and connect early models of region-specific instability to the human cancer genome.

## INTRODUCTION

Genome dynamics, encompassing the continuous and multifaceted changes within the genome (1, 2), underpin processes of adaptation, evolution, and disease progression (3). Dynamic mechanisms, such as DNA rearrangements, structural alterations, copy-number variations, epigenetic modifications, and activity of mobile genetic elements, drive genome plasticity (4–6), in part as an intricate interplay between the demands of genome conservation and the functional need for genome evolution (7, 8). One mechanism contributing to genome dynamics is genome rewiring/reconfiguration through the formation of extrachromosomal circular DNA (eccDNA; a term encompassing all classes of distinct extrachromosomal circular elements ecDNA, microDNA, spcDNA, etc. and collectively referred to here as the Circulome). Each overlapping class of circular DNA elements has emerged as a significant player in genome plasticity and/or functional diversification with far-reaching implications for health and disease (9–12).

The endogenous Circulome in eukaryote, primarily derived from linear chromosomal loci, exhibits specificity based on cell type, pathology, and developmental stage, contributing directly to somatic mosaicism. Biogenesis of extrachromosomal-circular DNAs impacts gene regulation, function, and copy-number variation. A prominent example of the connection between elevated levels of circular DNA and perturbation of key biological processes is oncogenic amplification driven by a subclass of eccDNA, ecDNA, known classically as double-minute chromosomes in their paried form. In addition, eccDNA mediates specific developmental programs, functional diversification, genome instability, and disease progression (12–14). Certain genomic loci are prone to inclusion among extrachromosomal circular DNAs, provoking the question of whether there may be hotspots of localized genomic plasticity (15–18).

A well-studied hotspot of chromosomal instability is the telomere, with telomere shortening known in some cases to extend beyond regular telomeric repeats (19–22). Moreover, McClintock described almost 100 years ago distal chromosomal regions in maize unique in their capacity to participate in genome recombination and dynamics (23–25). More recently, the subtelomeric position of COMMD5 near the end of chromosome 8 has been proposed to influence its expression and cancer-associated functions (26). Recent research has shed light on a functional role of extrachromosomal-circular DNA in telomere maintenance (27, 28). Circular DNA molecules carrying telomeric repeats have been shown to participate in telomere recombination (29–35), effectively counteracting telomere shortening and bypassing cellular senescence in cancer stem cells. This connection between extrachromosomal-circular DNA and telomere dynamics underscores a critical role of these elements in genome stability (36–39).

Here we use endogenous extrachromosomal circular DNA as a reporter of genome dynamics to investigate chromosome-end instability beyond telomeres/sub-telomeres. We find that extrachromosomal-circular DNAs are highly enriched in megabase-scale segments at select ends of specific chromosomes, pointing to increased dynamics at these distal regions. Significant enrichment of eccDNA extends well beyond the telomeric and subtelomeric regions, suggesting extensive chromosome erosion and genome “fragility,” elevating the potential for specific genes or regions to be rendered functionally aneuploid and/or incorporated into effective tumor-driving replicons. We find that this propensity for chromosome-end crisis is evident in both cancer cell lines and human cancer tissues, with varied intensities as a function of cell type, selective pressures, and pathology. Together, these findings reveal a previously unrecognized nexus of chromosome-end instability captured by a new class of Circulome elements. Our results illuminate a fundamental mechanism of genome remodeling with broad implications for cancer evolution, therapeutic vulnerability, and genome engineering.

## MATERIAL AND METHODS

### Cell lines and human tissues

For this work, we obtained multiple human cell lines and tissues for analysis.

i. Genomic DNA was extracted from a panel of cell lines: IMR90 and BJ (primary fetal lung fibroblasts), HeLa (cervical carcinoma), A549 (lung adenocarcinoma), U2OS and Saos2 (osteosarcoma). Two immortalized derivatives of IMR90 cells were also used: SW39 and SW26. Based on their telomere maintenance mechanisms, IMR90 and BJ are telomerase- and ALT-negative primary fibroblasts, HeLa and A549 are telomerase-positive cancer cell lines, whereas U2OS and Saos2 maintain telomeres via the alternative lengthening of telomeres (ALT) pathway. SW39 and SW26 are SV40-immortalized derivatives of IMR90 that utilize telomerase and ALT, respectively.
ii. For comparison, non-cancer datasets were obtained from fibroblasts and granulocytes from previous whole-genome sequencing and eccDNA studies of a male individual (13, 40); fibroblast cells were derived from a punch biopsy of healthy skin, while granulocytes, present at high levels due to myelofibrosis, were isolated from blood.
iii. Genomic DNA from a noncancerous reference cell line (Genome in a Bottle, GIAB) was included as a well-defined reference; this DNA was obtained from NIST (41).
iv. Ultra-high molecular weight (UHMW) genomic DNA was isolated by manual spooling from a glioblastoma cell line (GBM3094), which was directly derived from patient tissue.
v. A matched set of flash-frozen Adjacent Normal (AN) and Bladder Cancer (BC) tissues, resected from bladder cancer patients, was obtained from the UTSW Medical Center under IRB STU102014-008 and a material transfer agreement (MTA). Tissues were either pulverized using the COVARIS system or dissociated enzymatically into single cells following published protocols (42). While cell viability ranged from 30% to 80%, depending on sample quality, freezing/storage conditions, and dissociation protocol; samples had intact nuclei.

### Isolation of Genomic and Circular DNA

UHMW genomic DNA was isolated from dissociated AN and BC tissue samples by manual spooling (43–45). In parallel, single-cell suspensions were prepared and optimized for comprehensive isolation of extrachromosomal circular DNA directly from cells (13, 46).

To enable both enrichment and comprehensive profiling of the full eccDNA repertoire in a size- and sequence-agnostic manner, we employed multiple complementary methods to selectively deplete linear DNA and/or purify circular DNA based on the unique topological properties of circular DNA. Specifically, three approaches were used (**See Supplementary Fig. S1A**):

1. **Circulome-Seq V1.0 (CSI)** – an exonuclease-based protocol as previously described (13);
2. **Cesium chloride–ethidium bromide (CsCl-EtdBr) ultracentrifugation** – a classical biophysical method for topology-based DNA fractionation (9);
3. **Circulome-Seq V2.0 (CSII)** – a novel, dye-free, enzyme-free protocol enabling topological fractionation of DNA directly from complex biological mixtures including cells, hydrogel-embedded cell lysates, and ultra-high molecular weight DNA (47).

We note that none of these protocols rely on column-based isolation or rolling circle amplification (48–50), as these method have been shown to introduce difficult-to-control (uncontrollable) biases confounding distribution measurements for eccDNAs and characterization of differential repertoire of eccDNAs (48–50).

For enzyme-dependent enrichment, we selected CSI as the primary protocol due to its significant advantages over other enzymatic methods. Specifically, CSI avoids the aforementioned rolling circle amplification (RCA) and other pre-library amplification or selection steps, thereby preserving the native topological and sequence complexity of eccDNA, minimizing artefactual reads, and improving reproducibility (51). The CsCl-EtdBr method, while useful for isolating covalently closed circular DNAs based on topology, has a size limitation, with reduced efficiency for capturing circles larger than ∼100 Kb (52). Additionally, CsCl-EtdBr cannot recover nicked circles, which co-band with linear DNA. In contrast, CSI enriches for both nicked and covalently closed circles but, like all enzymatic approaches, remains limited in capturing large circles due to the intrinsic fragility of DNA, which is susceptible to shearing from handling, pipetting, extended enzymatic treatment, and other mechanical stresses (53, 54). To overcome the limitations of both approaches, CSII was developed, which efficiently isolates large circular DNA species with no apparent upper or lower size restriction and captures both nicked and covalently closed forms. Unlike enzyme-based protocols and CsCl-EtdBr gradients, CSII significantly reduces in-aqueous shearing, enabling recovery of fragile and large eccDNAs.

One potential explanation for the apparent enrichment of eccDNA at chromosome ends was that it reflected a technical artifact of the ExoV (RecBCD) digestion step rather than a true biological signal. Chi sequences (5′-GCTGGTGG-3′) act as regulatory hotspots for RecBCD: upon encountering a Chi site, the enzyme pauses, inactivates its 3′→5′ exonuclease activity, and begins generating 3′-ssDNA overhangs to initiate homologous recombination (55–58). Because Chi-induced pausing locally protects DNA from degradation, regions enriched in Chi sequences could, in principle, be preferentially retained during ExoV treatment and appear as spurious eccDNA peaks. While the use of exo-independent assessments of eccDNA distribution rules this out as the source of end enrichment, we nonetheless examined the genomic distribution of Chi sites relative to our eccDNA signal in exo treated samples and found no correlation between Chi sequence density and eccDNA-enriched regions.

A major obstacle in the eccDNA field has been the absence of standardized reference materials to establish a “ground truth” for quantitative evaluation of isolation and analysis workflows. To address this, we designed and implemented custom standards to benchmark our technologies across various biological backgrounds, including human samples. Our control sample contained a defined mixture of 70% linear lambda DNA (known sequence) and 30% synthetic circular DNAs (“eccDNA mimics”) of various sizes and unique sequences, with no overlap between the circular and linear sequences. This sample was sequenced on both Illumina and ONT platforms for validation. It was then processed using the CSII workflow to assess enrichment performance. For CSII enrichment, approximately 100 ng of DNA was subjected to dye-free, enzyme-free topological fractionation in dense chaotropic salt solution adjusted to pH 7.0. Samples were ultracentrifuged 3 h at 22°C. The upper fraction was recovered as the circular-DNA-enriched fraction, then desalted by dialysis, concentrated by vacuum centrifugation, and processed for downstream library preparation and sequencing. Reads were mapped to both lambda DNA and a reference set of the eccDNA mimics. We observed >70-fold enrichment of eccDNA mimics (calculated as the ratio of relative abundance post-enrichment versus input), and >95% removal of linear lambda DNA. The resulting eccDNA purity in the control sample exceeded 95% (**Supplementary Fig. S1B**).

### Library preparation and high-throughput sequencing

For all data presented in this study, we used Tn5 (Tagmentase; Illumina) to fragment both genomic DNA and eccDNA fractions (59), generating sequencing libraries compatible with both Illumina and ONT platforms. For each DNA sample, at least two independent tagmentation reactions were performed, and a minimum of two libraries were sequenced. Libraries were processed on Illumina’s MiSeq or NextSeq2K (P1 flow cell) for pooling and quality control. Initial next-generation sequencing (NGS) was performed using the NextSeq2K, NovaSeq6000, or NovaSeqX platforms. Batch effects and instrument-specific biases were systematically evaluated. Due to detectable biases associated with the NovaSeqX platform, we restricted subsequent sequencing to the NextSeq2K and NovaSeq6000. All reads generated were paired-end.

To assess technical robustness, we evaluated the reproducibility of our eccDNA enrichment and sequencing workflow. The protocol demonstrated >92% reproducibility, supporting its reliability for downstream analysis and distinguishing this workflow from alternatives that have lower reproducibility exhibiting technical variability across multiple datasets (48–50).

### Bioinformatic analysis

For characterization of both eccDNA and genomic DNA, paired-end FASTQ files were analyzed using a custom software pipeline composed of the following modules:

i. a k-mer–based mapper to align eccDNA reads to the reference genome; (ii) classification of reads into three categories—unique, focal repeats, and dispersed repeats—to reduce mapping artifacts and preserve repetitive content; (iii) high-sensitivity detection and evaluation of circular, deletion, gene-fusion, and insertion junctional reads (i.e., sequences that form novel junctions absent from the linear genome); (iv) post-analysis quality control to evaluate library quality and assign confidence scores to predicted variants; (vi) a minimum eccDNA enrichment calculator to evaluate enrichment relative to matched genomic DNA samples (in addition to other statistical approaches); (vii) a copy number estimation module based on spike- in standards and related metrics; and (viii) a visualization package for rendering and interpreting analysis outputs.

The visualization involves a two-dimensional plot where the vertical position of each segment (bin or gene) indicates the fold-enrichment in the corresponding uniquely mappable eccDNA fractions relative to unenriched (*i.e.*, total genomic) DNA, gDNA, with horizontal position indicating chromosome location (see Fig. 1). Enrichment of specific genomic regions in eccDNA samples (compared to total DNA) is thus quantified in terms of *y*-axis amplitude. For completeness of data display (but not used for the conclusions drawn here), we add a color scale to indicate a second, eccDNA-independent, enrichment metric: warmer colors here indicate areas where read counts in the total (unfractionated) DNA data are higher than expected based on assumptions of roughly equal capture of unique k-mers across the genome. The color metric compares different regions (from the same sample) that may have different capture efficiencies. Thus color values may be slightly skewed within a sample by region-specific base composition differences, Strong and consistent differences between biological samples are not subject to such issues, and we note that the values for stable (non-cancer) genomes are much “flatter” on both *y* axis and color scale, indicating that differences in both dimensions reflect properties of cell populations and not simply the underlying sequence.

**Fig. 1.**
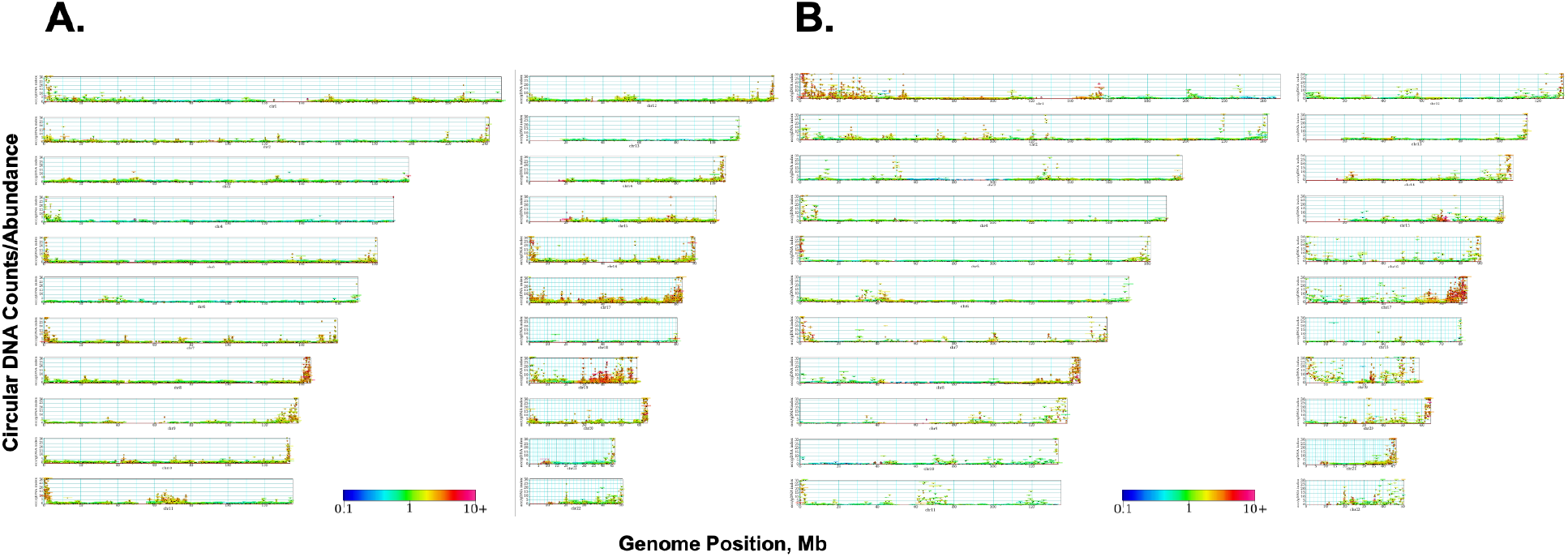
Whole-genome Circulome Maps. Enriched eccDNA signal relative to the corresponding gDNA for various cell lines. The *x* axis corresponds to chromosome coordinates. Abundance of eccDNA relative to total DNA of each 30-kbp bin of unique k-mers is plotted on *the y* axis and is capped at 30-fold. The color of the points is a function of calculated apparent amplification of each bin, with warmer colors indicating a higher level of copy number amplification. As examples, a full-scale red marker indicates that unique k-mers from the indicated bin are 10-fold more prevalent in sequencing data than expected based on fraction of total k-mers in the genome; a full-scale blue marker conversely would represent a bin where k-mers are 10 times less prevalent than expected. **A.** Circulome map for an immortalized cell line (BJ fibroblasts) **B.** Circulome map for an immortalized Saos2 (ALT), a human osteosarcoma cell line. [High-resolution plots-zooming shows full detail]. We note that only uniquely mapped reads are displayed. Genomic regions consisting entirely of tandem repeats, dispersed repeats or unassembled regions do not contribute to the signal shown. (*e.g.* telomeric sequences, Alu elements, p arms of chr 13, 14, 15)

### Sequencing-Based Characterization of Circular and Total DNA Pools

The sensitivity of existing bioinformatic tools is limited by the detection cutoff on the eccDNA size and the high background noise contributed by linear DNA sequences in WGS data. Here we demonstrate that this sensitivity barrier can be overcome by effective and unbiased enrichment of circular DNA targets using the CSI and the CSII technology. We generated WGS data from GBM3094 (GBM3094_gDNA) and sequencing data from the enriched circular DNA fractions using the CSI and CSII technologies (GBM3094_eccDNA). Further, we applied Amplicon Architect (AA) (60) on both the gDNA and eccDNA datasets. From the GBM3094_gDNA data, the AA algorithm predicted one ∼6 Mb circle with the following genes (EGFR, IKZF1, HSPB1, CDK6, and YWHAG). Using the GBM309_eccDNA datasets, AA predicts two circles (∼3 Mb and ∼7 Mb) harboring the following genes (CDC25A, USP4, GNAI2, AIMP2, EGFR, RHOA, CDK6, PMS2, RAC1, MST1R, RASSF1, PLXNB1, and IKZF1). Additionally, applying our eccDNA-specific informatic pipeline to characterize the repertoire of eccDNAs in this sample shows that AA fails to detect an abundant class of eccDNAs that are < 1Mb, along with failing to detect EDGE-eccDNAs. This highlights the complementary value of direct-isolation method, which can capture giant eccDNAs that informatic tools such as AA miss due to the low sensitivity.

### Statistical analysis and ML

Following the custom software pipeline to process the eccDNA and gDNA fastq files outlined in sections (i) through (vii) in the eccDNA isolation/enrichment, sequencing, and analysis section above, we generate the statistical and machine learning results following a process of:

a. First calculating the eccDNA enrichment metric, and amplification metric at 30Kbp segment sizes for both cell line and bladder cancer samples.
b. Create an ‘End only’ dataset, wherein we retain only those segments within 1Mpb of the p-arm, and q-arm for each chromosome.
c. Create an ‘Interior only’ dataset, wherein we retain only those segments that are at least 1Mbp distant from both the p-arm and q-arm of each chromosome.

For the ‘End only’ dataset, we train a stochastic gradient boosted tree classifier on end segments corresponding to each chromosome independently. For each chromosome, 50 train/test splits are randomly chosen, and the mean AUC is reported for each chromosome. This process is repeated 2 times, once for the amplification metric and once for the enrichment metric. For the ‘Interior only’ dataset, we follow the same process as just described, however for each chromosome, we sample a number of regions from the interior such that the dimensionality (number of features) matches the ‘End only’ dataset in order for a fair comparison to be made. We repeat this sampling process 50 times (completing a training process that involves 50 train test splits identically as described above), resulting in 50 mean AUC values for each chromosome, for each of our two metrics. We then report the means of these 50 AUC values per chromosome in a box and whiskers plot, showing means and quartiles of the AUCs with outliers.

## RESULTS

### Terminus-Proximal Chromosome Regions with Increased Genomic Plasticity are Revealed through eccDNA Isolation and Profiling

To define chromosomal regions that exhibit elevated probability of circular-DNA formation (and hence higher degree of plasticity) across pathologies and cell-types, we obtained multiple human cell lines and tissues for analysis. (*i*.) Genomic DNA was extracted from a panel of cell lines (see **Methods**); (*ii*.) comparable non-cancer datasets were obtained from fibroblasts and granulocytes from previous whole-genome sequencing and eccDNA studies of a male individual(13, 40); (*iii*.) genomic DNA from a noncancerous reference cell line (Genome in a Bottle, GIAB), which is a well-defined reference; (*iv*.) a matched set of flash-frozen Adjacent Normal (AN) and Bladder Cancer (BC) tissues, resected from bladder cancer patients.

To enable both enrichment and comprehensive profiling of the full eccDNA repertoire (the Circulome) in a size-and sequence-agnostic manner, we employed a set of robust biophysical strategies that selectively deplete linear DNA and/or purify circular DNA by exploiting the unique topological properties of circular molecules. Specifically, three independent approaches were used and followed by bioinformatic analysis (see **Methods** and **supplementary Fig. S1**). The first approach is Circulome-Seq (CSI), an exonuclease-dependent enrichment of circular DNA that we previously described in (13). The second approach is CsCl-EB gradients, an enzyme-free isolation method that isolates DNA based on topology, which is the gold standard for isolating covalently-closed circular DNA (9). The third method is a new method we developed for enriching eccDNAs in an enzyme-independent manner (Circulome-Seq 2.0) (47), which improves on CSI’s capture efficiency of eccDNAs >500 kb.

Across all analyzed cell lines, we observed an enrichment of eccDNA originating from specific chromosomal ends. This pattern reflects a form of chromosome-end dynamics characterized by a significantly higher probability of circularization at distal chromosomal regions (**Fig. 1**). Notably, this effect can extends megabases inward from chromosome termini into gene-containing regions, well beyond canonical telomeric and subtelomeric domains, indicating a form of chromosome-end instability that is mechanistically distinct from telomere shortening or simple linear-end erosion. It is worth noting that the GIAB cell line and normal fibroblasts exhibit an attenuated chromosome-end signal relative to the derived cancer-cell lines (**Fig. 2A**).

**Fig. 2.**
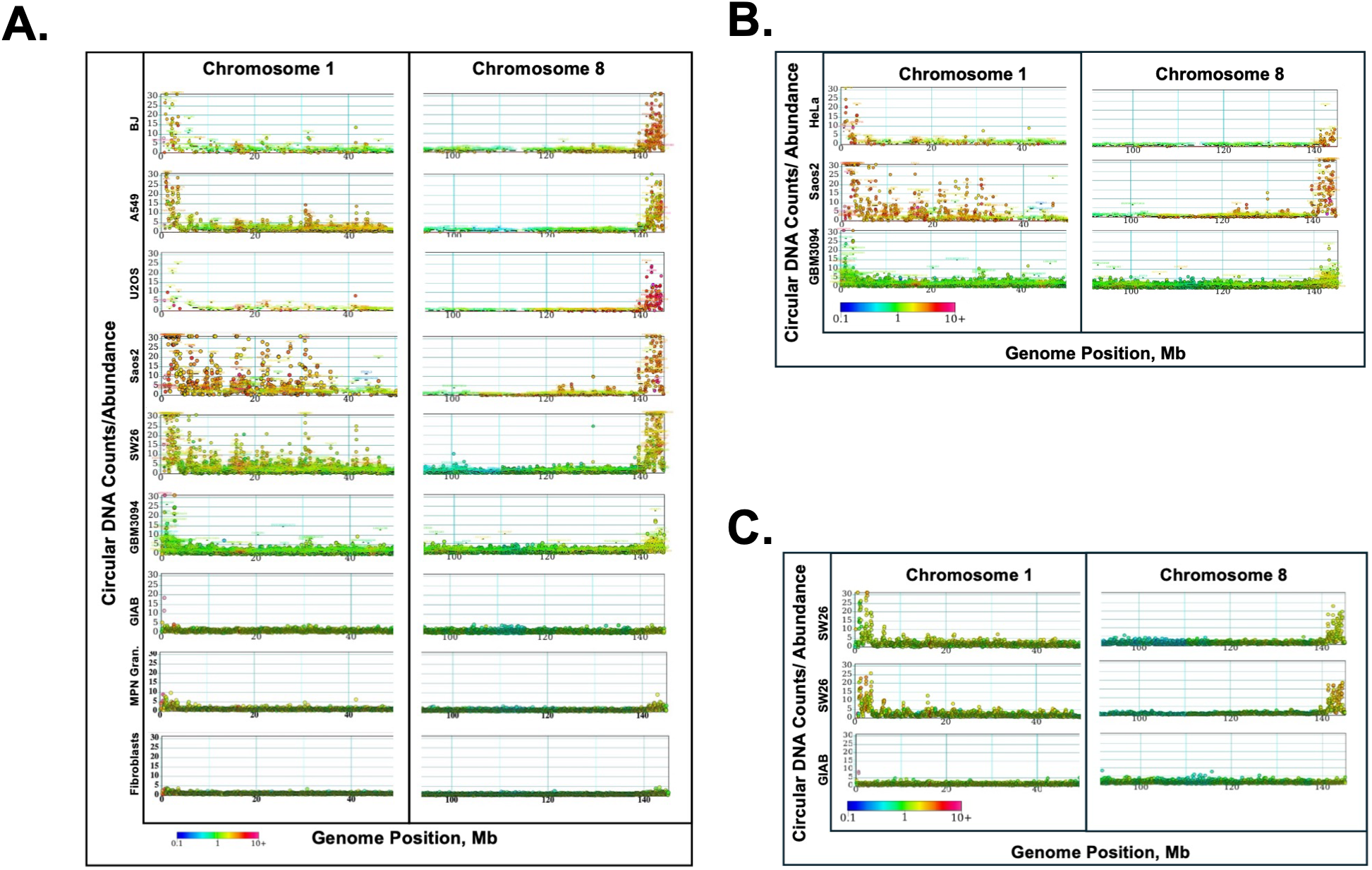
Assessments of robustness for terminal eccDNA enrichment in specific cell populations. **A.** ExoV treatment-enriched eccDNA signal (CSI protocol) showing positional enrichment of eccDNA relative to the corresponding gDNA for various cell lines. For clarity, we highlight just two regions of the genome here, the terminal ends of chromosomes 1 and 8. **B.** Dual-enzyme validation of ExoV analysis. The eccDNA fractions from ExoV (CSI) treatment were further processed with lambda-Exo to reduce any putative residual linear DNA that might have survived digestion with ExoV. We note that the eccDNAs mapping to the telomere-proximal coding regions remain enriched after additional treatment with the dsDNA-specific lambda exonuclease [full maps can be found in supplementary material]. **C.** Nuclease-independent fractionation of eccDNA using CsCl-EtdBr equilibrium density-gradient ultracentrifugation. Here CsCl-EtdBr fractionation of genomic DNA was used to isolate covalently-closed circular DNAs from select cell lines. On the top panel, we show cancer cell line SW26 eccDNA enrichment from the bottom band of the gradient relative to the top fraction of the gradient, which contains both linear DNA and nicked-circular DNAs. The middle panel shows the eccDNA enrichment from the bottom band of the gradient relative to total corresponding genomic DNA (linear and circular). Note the reduction in overall enrichment signal. The bottom graph shows the eccDNA enrichment in reference cell line GIAB from the gradient bottom band relative to the corresponding total genomic DNA. The color of the plot is a function of calculated apparent amplification total DNA for each bin, with warmer colors indicating a higher level of amplification.

### Observed eccDNA Boundary Enrichment is Robust to Isolation, Computational, and Statistical Methods

To further examine the observed enrichment of eccDNA at chromosome ends, we investigated whether this positional pattern was shared between fundamentally different eccDNA isolation strategies. Because ExoV–based enrichment could, in principle, introduce bias through incomplete digestion of linear DNA, we first evaluated eccDNA enrichment using an ExoV-independent approach. Specifically, we applied a modified Circulome-Seq workflow based on cesium chloride–ethidium bromide (CsCl–EtdBr) density-gradient ultracentrifugation to, which separates DNA molecules based on topology rather than enzymatic digestion, to SW26 cell line. Sequencing of the covalently closed-circular DNA fraction (the bottom band of the CsCl-EtdBr gradient) revealed a chromosome-end enrichment pattern comparable to that observed following ExoV-based enrichment. In contrast, this enrichment pattern was not observed in the fraction containing predominantly linear DNA with some nicked circles (top band), confirming that the signal is associated with circular-DNA topology. To further exclude the possibility that residual linear DNA surviving ExoV treatment contributes to the observed enrichment, we subjected ExoV-treated samples to an additional round of treatment with a different exonuclease, lambda-exonuclease digestion, which selectively degrades double-stranded linear DNA via a distinct enzymatic mechanism. The persistence of chromosome-end eccDNA enrichment following this additional treatment demonstrates that the signal is not dependent on a single nuclease or isolation strategy. Together, these results show that terminal eccDNA enrichment is reproducible across orthogonal enrichment methods and is therefore unlikely to arise from isolation-dependent or computational artifacts (**Figs. 2B** and **2C**). We also show a high reproducibility of enrichment technical replicates (**supplementary Fig. S2**).

Computationally, we employed two approaches: (i) conventional alignment and counting using BWA (61, 62) and (ii) a k-mer–based approaches (63, 64) to map and characterize both gDNA and eccDNA reads (**See Methods**). The k-mer strategy is particularly well-suited to eccDNA analysis: it is robust to repetitive structures, the circular junctions, and structural rearrangements that confound linear read alignment. In either case, eccDNA enrichment (defined as the ratio of the normalized read coverage of a given bin (or gene) in the eccDNA fraction to its normalized coverage in the matched gDNA fraction) is reproducible (**supplementary Fig. S3**).

With eccDNA enrichment at chromosome ends is apparent from the distribution plots (**Figs. 1** and **2**), we sought to quantify the extent to which enrichment depends on position relative to chromosomal boundaries. For this statistical analysis, we refer to chromosome ‘boundaries’ as genomic regions at the edges of large chromosomal domains, including both p- and q-arm termini (para-telomeric) as well as centromere-adjacent regions (peri-centromeric), and contrast these with interior chromosome regions. Resolving large-scale structure in eccDNA enrichment patterns across all chromosomes is facilitated by enrichment aggregation within 1-Mbp segments for all chromosomes end to end (p:q arm).

We constructed the empirical cumulative distribution (ECDF) (65) of eccDNA enrichment level for each chromosome enabling quantification of observational likelihoods for eccDNA enrichment levels at chromosome-wide resolution. Applied to GBM and stage-1 bladder-cancer tissue, eccDNA-enrichment levels in the upper-most quantiles are dominated by segments adjacent to chromosome boundaries (**Figs. 3A** and **3B**). Further, high levels of eccDNA enrichment are not stochastically dispersed, but rather exhibit “volatility clustering”, in which segments of high eccDNA enrichment are more likely to be adjacent to other segments of high enrichment. This phenomenon is formally characterized by measures of significant spatial autocorrelation of eccDNA enrichment along the extent of a chromosome (**Figs. 3C** and **3D**), and is analogous to a phenomenon in financial time-series data wherein the volatility of a financial instrument exhibits temporal clustering (conditional heteroskedasticity) (66).

**Fig 3.**
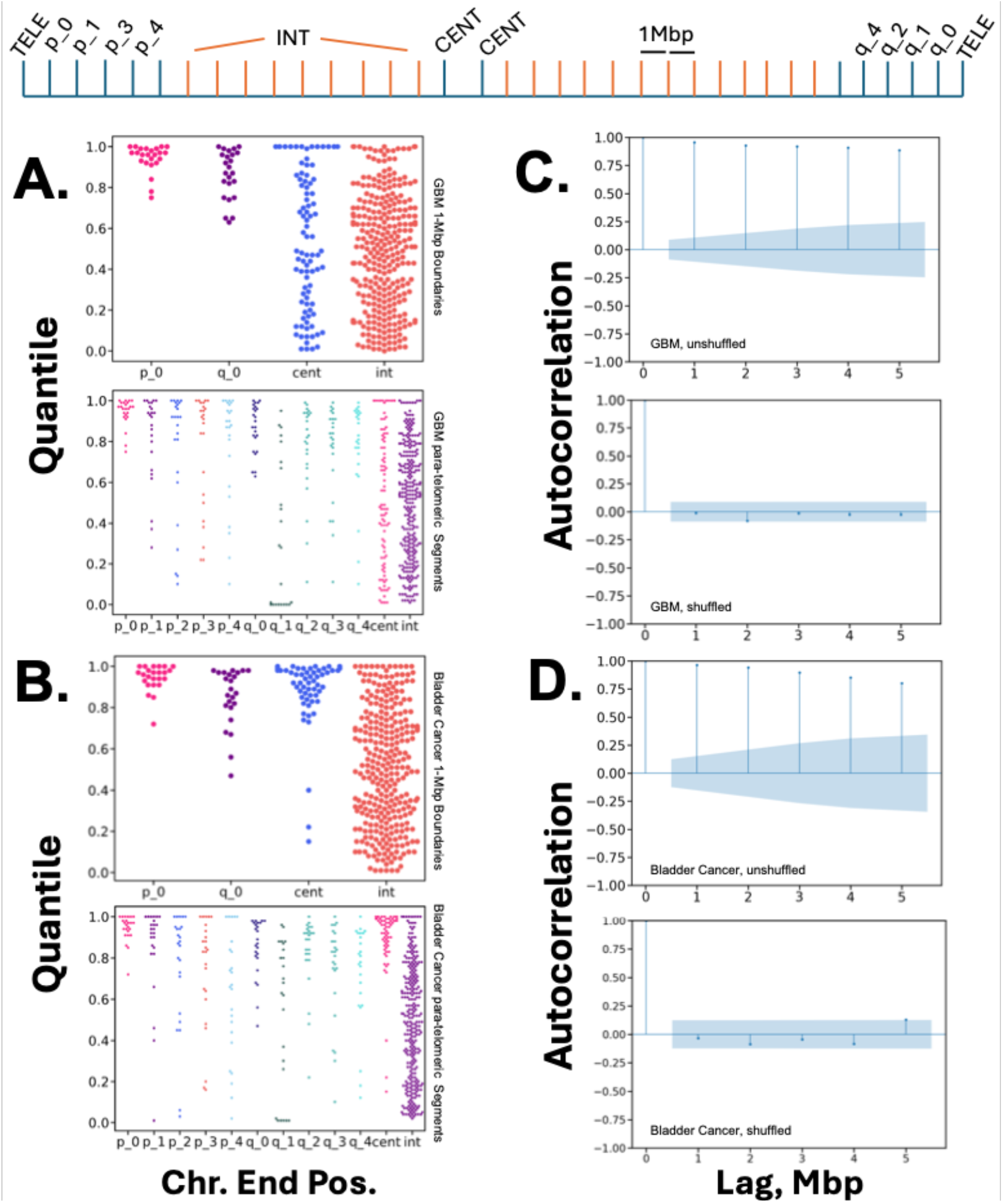
Chromosomal End Enrichment Clustering of eccDNA. The schematic at top shows the spatial arrangement of named X axis constituents in A and B. p_0 is the 1Mbp chromosome segment closest to the p-arm telomeric end, and q_0 similarly is the segment adjacent to the q-arm telomeric end. Orange segments (INT) represent chromosomal segments interior of p,q named segments and the centromere adjacent segments (CENT). Four panels are shown: GBM1 (A, C) and a Bladder Cancer Stage 1 sample (B, D). Panels **A** and **B** show percentile plots from the empirical cumulative distribution functions (ECDFs) constructed using eccDNA enrichment scores defined for select 1-Mbp chromosomal segments from each chromosome. The Y axis shows the ordinal percentile of the magnitude of eccDNA enrichment for each selected segment category (e.g., a value of 1 indicates the highest level of enrichment seen for any segments genomewide, 0.5 represents median enrichment level). For the X axis, segments p_0 and q_0 correspond to the para-telomeric regions (p-arm end of chromosome and q-arm end of chromosome), cent denotes the centromeric-adjacent region, and int corresponds to those regions of each chromosome that are neither para-telomeric, nor peri-centromeric (*i.e.* interior regions of the chromosome). The top subplots from **A** and **B** show ECDF values for all chromosomes, comparing p_0, and q_0 with the centromeric and interior regions. The bottom subplot shows the same data but include segments increasingly distant from the para-telomeric regions (p_1 is interior of p_0, p_2 interior of p_1, etc.). Panels **C** and **D** show the autocorrelation function calculated using a series of 1Mbp segments spanning chromosome 1 from GBM1 (**C**), and the Bladder Cancer Stage 1 sample(**D**), ordered from the start of each chromosome (para-telomeric p-arm) through terminus (para-telomeric q-arm). The Y axis denotes correlation values between adjacent chromosomal segments (a value of 1 indicates perfect correlation, and 0 indicates no correlation). The X axis denotes ‘Lags’, which are 1Mpb chromosomal segments immediately following one another in the spatial order of the chromosome from p-arm to q-arm (Lag 0 is always perfectly correlated with itself). For example, if p_0 is Lag 0, then p_1 is Lag_1, and p_2 is Lag 2, etc. The top subplots in each of **C** and **D** show the autocorrelation values (averaged, calculated for all windows of 1 to 5Mbp along the full span of the chromosome) for the true ordering of segments along the chromosome. Each point represents the autocorrelation value for that lag, and the blue bands represent an uncertainty region within which autocorrelation would not be significant (the stem does not represent a quantitative value, serving only as a visual aid to anchor each autocorrelation value to the base and uncertainty/insignificance region). These plots show that correlation with Lag 0 remains strong for at least 5 1Mbp segments following Lag 0 (Lags 1-5), demonstrating spatial correlation of eccDNA enrichment values. The bottom subplots in C and D each repeat the same calculation, however we first randomly shuffle the segment ordering, thus the ordering of 1Mpb segments no longer reflects their true position along the span of the chromosome. After this shuffling, we observe no autocorrelation at any of the lags, confirming that there is significant ‘spatial clustering’ of eccDNA enrichment values for segments ordered according to their true positions along the chromosome.

### Structural rearrangements are Enriched in Megabase-Scale Regions Proximal to Chromosomal Boundaries

From the observation that chromosome ends are enriched in eccDNA, we sought to understand to what degree the incidence of structural rearrangements at chromosomal boundary regions would show similar enrichment. We investigated 3 junction types: Circle, Insertion, and Deletion at 10-Mbp resolution, aggregating observations of each class within each 10-Mbp window constructed from p-arm to q-arm. We accumulated incidences for each class within each 10-Mbp bin and plotted these incidences as bars for each chromosome from p-arm (leftmost) to q-arm (rightmost) (**Fig. 4)**. The majority of chromosomes have peak incidences at the chromosomal end regions indicating that these boundary regions also show a similar enrichment pattern. These findings support the hypothesis that EDGE is reporting on regions exhibiting multiple modes of instability; effectively a higher accumulation of genomic rearrangements in these chromosomal regions.

**Fig 4.**
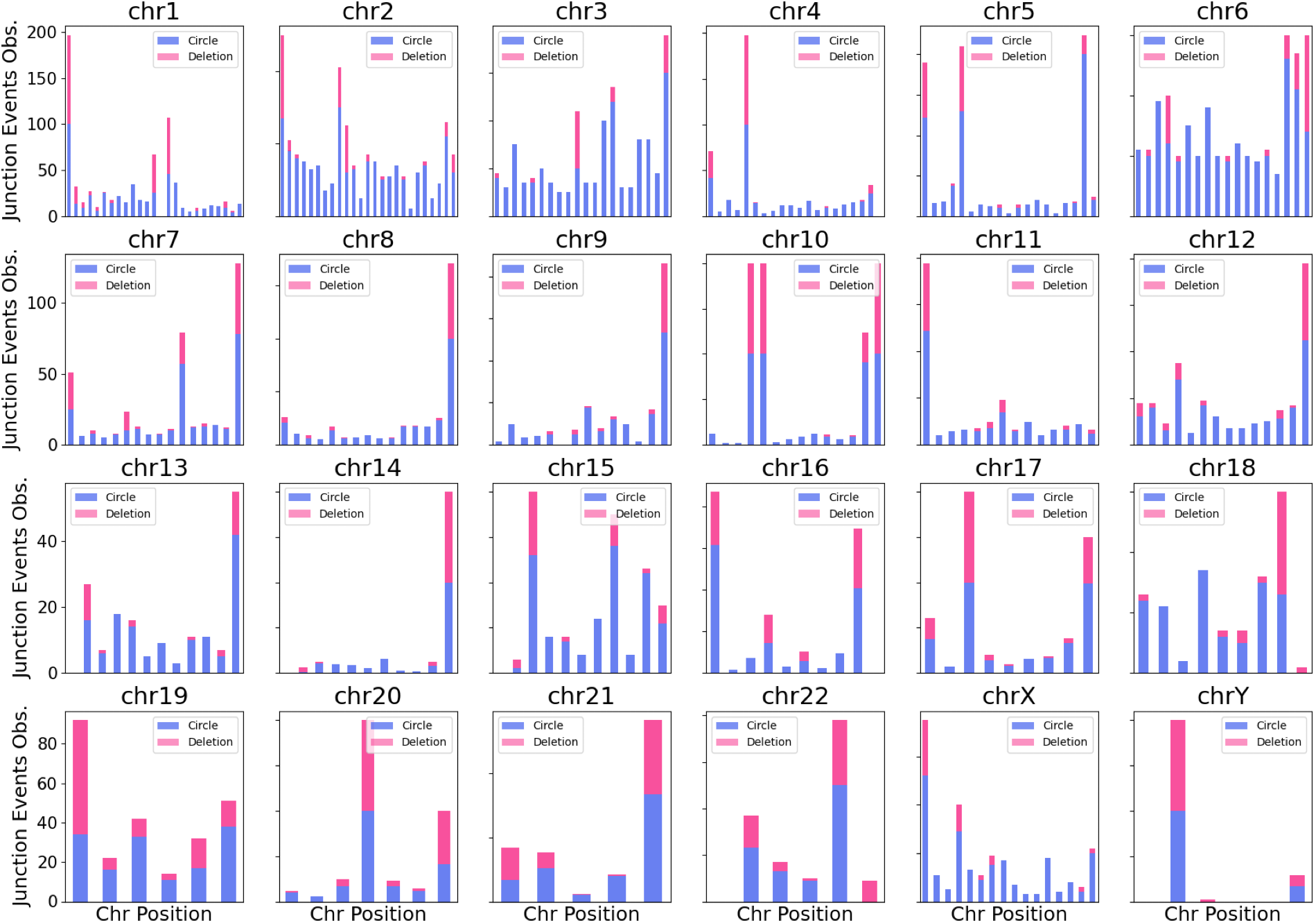
Junction Events by Chromosomal Location. The incidence of two different Junction event classes as a function of chromosomal position: Deletion and Circular junction counts are plotted for 10-Mbp segments of each chromosome for the Saos2 cell line. The Y axis represents the counts for each Junction type observed at a particular 10-Mbp region. These counts are plotted along the X-axis, which represents chromosome coordinates. The resulting stacked bar plots for each chromosome indicate areas of overlap and exceedance in the count values for the two Junction Event Types. Events accumulated within 10-Mbp regions are displayed along the x-axis of each chromosome, from p-arm (leftmost) to q-arm (rightmost), capped at an incidence of 100 observations for the representative dataset used in this analysis.

### eccDNA Enrichment at Chromosomal Ends Enables Accurate Classification of Cancer Status for Marched Normal and Tumor Cancer Tissues

To determine if the eccDNA-specific end effect is a feature exclusive to cell lines, we investigated chromosome-end dynamics in human bladder tissues of varying pathologies. For this work, we investigated 40 matched bladder-cancer (BC) and adjacent-normal (AN) tissues (paired samples from 20 patients). These tissue samples were subjected to pathological characterization. Pathological classification of “Normal” vs. “Cancer”, including % tumor in the cancer samples was performed and reviewed by a genitourinary pathologist at an NCI designated cancer center (UTSW).

Analysis of these data revealed a striking enrichment of eccDNA at chromosomal boundary regions (**Fig. 5B**). Enrichment was also observed at non-telomeric chromosome termini and centromere-adjacent regions within chromosome interiors (**Fig. 5B**). These patterns were consistent across multiple patients and were not limited to a single pathology subgroup in this cohort.

**Fig 5.**
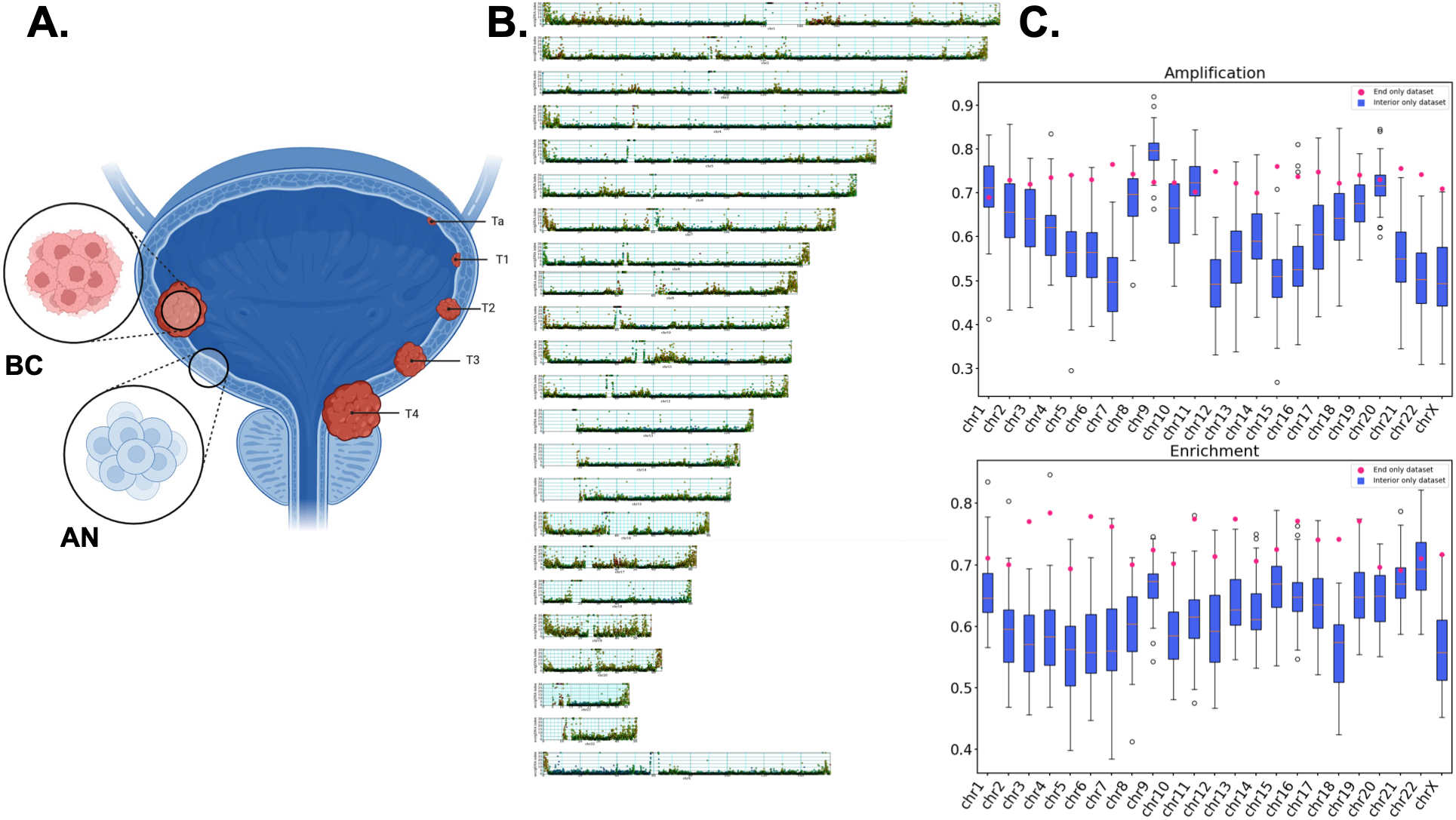
eccDNA end enrichment distinguishes bladder cancer from adjacent normal tissue. **A.** We processed and analyzed matched bladder cancer tissue and adjacent normal from 20 bladder cancer patients (various stages of disease) who have gone through complete bladder resection. **B.** Whole-genome Circulome map for a representative bladder cancer (BC) tissue sample, showing enrichment of eccDNA near chromosomal boundary regions. **C.** Classification performance per chromosome using amplification features (top), and enrichment features (bottom). Amplification features are copy number amplifications calculated solely from linear genomic DNA. Enrichment features are calculated from a normalized ratio of eccDNA copy number over linear genomic DNA copy number, and thus represent a robust value for eccDNA enrichment. For the ‘Interior only’ dataset, given that its feature dimensionality is much larger than the ‘Ends only’ dataset, to make a fair comparison we sample the same number of features in order that the feature dimensionality matches the ‘End only’ dataset. We repeat this feature sampling process 50 times (keeping the same number of 50 70/30 train/test splits within each repetition) and report the resulting AUC values for each repetition as a box and whiskers plot in blue. The resulting AUC for the ‘Ends only’ dataset is plotted as a red point. We see in the amplification results (top) that with only a few exceptions, the AUCs of the ‘End only’ dataset dominate those from the interior—with many of the mean AUCs of the ‘Interior only’ dataset being at, or very near .5, indicating there is no information gained about the class assignment AN/BC. The enrichment results (bottom) show that for all chromosomes the AUC values for the ‘End only’ dataset are higher than the means from the ‘Interior only’ dataset. Taken together, we demonstrate that for all chromosomes, for both amplification and enrichment, the ‘End only’ dataset is always informative, and that in most cases, across both amplification and enrichment, it yields AUC values significantly above those obtained from a dataset comprised of only interior regions of each chromosome.

We then ask the question of whether or not eccDNA enrichment at chromosomal boundaries has classification utility using both classical (CNV/Amplification) features and enrichment features constructed from eccDNA enrichment (**Fig. 5C**). For this analysis we create an ‘End only’ dataset defined as chromosomal regions within 1Mbp of chromosome ends, and an ‘Interior only’ dataset defined as those regions not within 1Mbp of chromosome ends. We train a stochastic gradient-boosted tree classifier (67) on these two datasets (see **Methods**) for two feature classes: amplification/CNV (**Fig. 5C** top) and eccDNA enrichment (**Fig. 5C** bottom), for each chromosome. To enable a fair comparison given the greater feature dimensionality (number of 1Mbp regions) of the ‘Interior only’ dataset, we repeat our training process (70/30 train/test split for 50 splits) 50 times, sampling interior regions at each trial to match the dimensionality of the ‘End only’ dataset. We report the means of each trial’s AUC as a blue bar for each chromosome’s box and whiskers plot. We report the mean AUC of the ‘Ends only’ dataset as a red point in each panel. The amplification panel (**Fig. 5C** top) shows the mean for the ‘Interior only’ dataset at ∼ 0.5 (chance) for several chromosomes, indicating amplification/CNV for the interior regions of these chromosomes is uninformative with respect to distinguishing AN/BC. This is not the case for the ‘End only’ dataset, where for each chromosome the mean AUC is significantly above 0.5. We note though that chromosome 9 in the amplification panel appears to be a positive outlier (mean AUC 0.8), indicating a potentially significant locus. The eccDNA enrichment panel (**Fig. 5C** bottom) shows the ‘End only’ dataset exceeding the mean AUC of the interior only dataset for every chromosome, demonstrating that eccDNA enrichment values at chromosomal boundaries are always strongly informational with respect to distinguishing AN/BC.

## DISCUSSION

### Erosion at Distal Genomic Ends

Our study reveals a dimension of cancer-genome dynamics centered on the formation of circular DNAs from chromosome boundaries/ends, a process we term *Erosion at Distal Genomic Ends [EDGE]*. The striking nature of this signal led us to undertake multiple independent lines of investigation to evaluate and understand this signal. Using rigorous biophysical approaches for direct isolation and characterization of eccDNAs, we demonstrated that a new class of eccDNAs arise preferentially (and asymmetrically) from specific telomere-proximal coding and non-coding regions in a manner that reflects aspects of cell and tissue states. These findings mark a shift in our understanding of chromosome instability: whereas local telomere erosion has long been associated with genomic dysfunction, we show that extrachromosomal-circular DNAs that originate from chromosomal sites far from genic/non-genic boundaries can report on a broader and previously uncharacterized form of chromosome-end degradation and complex structural variation, with a scale that penetrates well beyond the telomeric and subtelomeric regions.

This work builds upon prior foundational studies: (i) McClintock’s work that established chromosome ends and distal regions are not passive termini but uniquely capable of breakage, fusion, recombination, and structural remodeling (23–25, 68) (ii) Detailed studies of telomere-driven chromosome instability (69–72) (iii) and evidence that the Circulome (a term that encompasses many distinct eccDNA classes (12, 13); double minutes (73–75) a.k.a ecDNA (15–17, 76), microDNA (77–79), spcDNA (10, 80), etc.) can harbor and mediate the amplification of genes (*e.g.* double minutes/ecDNA), contributing to tumor evolution and therapy resistance (81–83). We extend this paradigm by identifying a new class of eccDNAs that are not only vehicles of oncogenic amplification but also act as high-resolution reporters of spatial genome instability. Unlike traditional whole-genome-based approaches for indirect extrachromosomal-circular DNA detection, direct eccDNA isolation and analysis can capture the structural and copy-number landscape of eccDNAs with high sensitivity. Notably, we identified a process in which eccDNAs form asymmetrically at the p and q arms of chromosomes, supporting the hypothesis that eccDNA biogenesis is spatially non-random. In this landscape we see the shadow of a cancer’s effect on the genome (84). Additionally, this work extends current models of cancer-chromosome dynamics. Indeed, among the cell lines investigated here, the osteosarcoma cell line Saos2 showed the most pronounced EDGE signal. This high level of genomic instability is consistent with previously reported high levels of genomic rearrangements in this model (85). However, our work also reveals EDGE-mediated chromosome-end dynamics as an additional and spatially defined dimension of this model’s genomic instability and dysregulation.

### Spatial clustering of highly enriched eccDNAs at chromosome ends

In addition to these findings, our analysis uncovered a striking *spatial clustering* of highly enriched eccDNAs, whereby regions of high enrichment tend to occur in contiguous chromosomal neighborhoods. This autocorrelation is reminiscent of “volatility clustering” observed in time-series financial models such as ARCH, where large fluctuations are temporally grouped (86, 87). In eccDNA distributions, the clustering occurs *spatially* along the chromosome, particularly at its terminal regions. These patterns suggest that eccDNA formation is not a purely stochastic process but one that reflects coordinated local chromatin states, chromosome accessibility, or mitotic mechanics (25). Other features are spatially enriched at chromosome ends, including repetitive elements (68), GC isochores (88), and hyperdivergent regions (89), and they exhibit late replication timing (90). The clustering of such features may likewise intertwine with additional cooperative biological mechanisms, potentially including epigenetic remodeling, e.g., focal histone modifications, DNA hypomethylation, or nuclear architecture constraints, that collectively predispose specific chromosome ends to structural fragility, including DNA cyclization. The convergence of elevated eccDNA formation and spatial clustering may represent a shared chromatin environment that facilitates coordinated excision and circle formation from neighboring loci.

### Possible mechanisms of EDGE

The molecular mechanisms underlying the boundary-specific enrichment of EDGE-associated eccDNAs remain to be determined. Several features of distal chromosome regions provide plausible explanations for this process. (15, 91, 71, 92). One possibility involves replication stress at subtelomeric and peri-terminal regions, which are known to experience replication challenges and may predispose these regions to fork collapse and DNA excision (24, 79, 93). Furthermore, the disruption of telomere–interstitial loops, often anchored by TRF2 and lamin A/C, could potentially destabilize adjacent genomic regions, introducing topological strain and increasing susceptibility to breakage (94–96). Consistent with this possibility, telomere position effect over long distances (TPE-OLD) has been proposed to modulate the expression of COMMD5, a subtelomeric gene near the end of chromosome 8 whose locus is recurrently affected by copy-number alterations in cancer (26). This work supports the broader principle that chromosome-terminal architecture can have biologically consequential effects on nearby gene regulation and provides a relevant context for examining local eccDNA dynamics at chromosome ends. In cancers, specifically in ALT-positive cells, repair-pathway preferences often shift from canonical non-homologous end joining (NHEJ) and homologous recombination (HR) toward microhomology-mediated end joining (MMEJ) (97, 98); this bias could favor excision-based repair outcomes that generate circular-DNA intermediates (99, 100). Notably, recent studies have also implicated epigenetic states as critical mediators of chromosomal fragility and cooperative genomic dynamics (101, 102). For instance, focal loss of heterochromatic marks and the redistribution of histone modifications, such as H3K9me3, H3K27ac, and H4K16ac, can render chromatin more accessible and permissive to excision-based remodeling (103). These epigenetic transitions may facilitate not only local circle formation but also spatial clustering of eccDNA production through coordinated chromatin states along adjacent genomic blocks, potentially contributing to the spread of instability from terminal regions into adjacent chromosome domains (104). Together, these observations suggest several possible models for why distal chromosome regions may be particularly susceptible to eccDNA formation, previous work has linked BFB to linear amplifications (105, 106), however, BFB and similar mechanisms as possible models of a class of extrachromosomal circular DNA amplification remain active area of research (75, 107). In any case, it is striking to observe the degree of enrichment of circular DNAs derived from chromosome ends.

### EDGE as a reporter of increased genomic instability

The enrichment of eccDNA at chromosome ends could in principle be explained by two non-exclusive models: (i) elevated rates of eccDNA formation at these loci, reflecting heightened genomic plasticity/instability, or (ii) preferential post-formation stability of chromosome-end-derived circles, possibly conferred by specific sequence content, specialized chromatin states, or proximity to telomeric structures. We acknowledge that the present study does not directly measure eccDNA formation kinetics or turnover, and our data therefore cannot, in isolation, distinguish a rate-of-formation effect from a stability/selective-retention effect. In either models, it is striking that the ends of chromosomes have a higher access to topological and copy-number diversity, leading to unique capabilities, roles, and responsibilities of these regions in normal and cancer tissue.

Nevertheless, several lines of evidence lead us to favor the instability/plasticity model. First, to our knowledge no specific sequence motif or structural feature is uniquely shared among chromosome boundaries and a subset of terminal regions yet absent from internal chromosomal regions, that would plausibly confer differential in-vivo nuclease resistance or segregational advantage to terminal-derived circles. Second, and more directly, we observe that independent measures of genomic instability (somatic deletions and structural variants) are enriched at the same chromosome-boundary regions that give rise to EDGE eccDNAs. This parallel enrichment is difficult to reconcile with a pure stability model, which would predict terminal eccDNA enrichment without a corresponding enrichment in linear-genome rearrangements. The convergence of these two orthogonal signals at the same loci is more parsimoniously explained by elevated local instability that manifests both as circular byproducts (eccDNA) and as linear genomic alterations (deletions, SVs). We therefore interpret EDGE as a readout of genomic plasticity at chromosome ends rather than as evidence of differential eccDNA persistence.

A related question is whether phenotypic selection in cancer cell lines may shape the Circulome. eccDNA carrying chromosome-end sequences may be retained in tumor populations if they confer fitness advantages, for example, by harboring regulatory elements, oncogenic loci, or sequences buffering against telomere attrition, however, the co-localization of EDGE peaks with somatic deletions and SVs argues against selection as the sole driver. The most parsimonious interpretation of the combined data is that chromosome ends are intrinsically more plastic genomic compartments, with both circular and linear instability arising from this shared underlying property. Selection in cancer would certainly act on these elements.

### Conclusions: context, limitations, and perspectives

We show that EDGE-associated eccDNA populations provide a quantitative readout of chromosome instability. Notably, EDGE eccDNAs are enriched in cancer samples examined in this work, highlighting their potential value as a sensitive biomarker of chromosome-end instability.

EDGE is distinct from classical aneuploidy, which involves stable gains or losses of whole chromosomes or chromosome arms (108, 109). Instead, EDGE is characterized by the enrichment of eccDNAs and/or copy number modulation of specific loci, as opposed to alterations of whole-chromosome copy number. EDGE is also distinct from classical telomere erosion, which hinges very locally on telomeric-sequence integrity and function (72, 107). In contrast, EDGE is associated with the enrichment of eccDNAs originating from regions extending megabases beyond telomeric boundaries, indicating a broader form of chromosome-end instability. Finally, EDGE is further distinct from ecDNA or double-minute elements, which exclusively harbor amplified oncogenes (e.g. EGFR, MYC, etc.) and hijacked regulatory elements (107, 110). In contrast, EDGE eccDNAs are enriched for genic, noncoding, and regulatory elements that map to chromosome boundaries on the linear chromosomes. Thus, EDGE eccDNA elements embed additional spatial information regarding their origin.

Although multiple genome instability pathways such as microsatellite instability (MSI), generic DNA-damage responses, or genetic background may contribute to or coexist with EDGE (111–113), EDGE itself is best understood as a spatial pattern of chromosome-end instability with eccDNA as its readout, rather than reflecting any single molecular mechanism. This work highlights EDGE-eccDNAs as a reproducible and informative marker of chromosome-end instability across diverse cellular contexts.

EDGE eccDNAs are unlikely to reflect fixed genomic endpoints, but rather seem to represent dynamic products of ongoing chromosome-end instability. We postulate that the fate of these products will depend on a complex interplay between the overall genetic background and selective pressures and may not always resolve into stable copy-number variants or gene loss. This process, however, might generate precursors that, with selective pressures, can yield a subset of stable extrachromosomal structural and copy-number alterations.

The analyses in this work provide several key capabilities: (*i.*) the ability to reproducibly isolate and analyze eccDNAs across a wide size and sequence range; (*ii*.) identification of chromosome-end-specific eccDNA enrichment and clustering; and (*iii.*) integration of these features into a machine-learning pipeline capable of accurate classification. Collectively, these findings advance eccDNA profiling as a potential tool in precision oncology. Notably, EDGE eccDNAs, as a distinct class of eccDNAs, may identify tumors with selective dependencies generated by chronic chromosome-end instability, potentially relying on compensatory pathways such as ATR/CHK1, WEE1, and POLQ to buffer genomic stress (114–118). This raises the possibility that EDGE-positive tumors could be preferentially sensitive to targeted inhibition of these DNA damage-response pathways, warranting further functional validation. Based on these promising results, it will be of substantial value to examine expanded sample cohorts. Our current models were trained and tested on datasets with limited sample size but achieved consistent performance through careful distributional modeling and random-permutation testing. Future work would optimally entail a Circulome Atlas database with integration of additional features such as circular-DNA methylation signatures, transcriptional capacity, and size distributions to further enhance biomarker resolution, target discovery, and therapeutic-prediction power.

## AUTHOR CONTRIBUTIONS

MJS and AZF originated the project. MJS supervised the project. MJS, AZF, and SDL directed the project. ZL and MJS and SDL did the experimental work. GR, MJS, AZF, MJM performed informatics analyses. GR performed ML and relevant analyses. Visualization was done by GR, ZL, MJM, AAS, AZF and MJS. Sample acquisition was done by YL, PCS, JS, AZF, and MJS. Writing of the original draft was done by GR, ZL, SDL, AZF, and MJS. All authors contributed to editing. Funding acquisition by SDL, AZF, and MJS.

## ACKNOWLEDGEMENTS

We thank Mark Ross, Sarah Harris, Nelson Hall, and Emily Greenwald for discussions. We thank Ellen Hong and Kavita Chalasani for providing materials that were used for this work. MS, GR, and SDL are affiliated with Phinomics, Inc. MS. and GR. are co-inventors on a patent covering the technology described in this work and its applications in biomarker and therapeutic target discovery. PS is now affiliated with Amgen. MJM is now affiliated with U. of Chicago and consults for Phinomics. YL consults for Photocure, Astra-Zeneca, Merck, Fergene, Nucleix, Ambu, Seattle Genetics, virtuoso surgical, Stimit, Urogen, Vessi medical, CAPs medical, Nonagen, Aura Biosciences, Inc., Convergent Genomics, Pacific Edge, Pfizer, Phinomics Inc, CG oncology, Uroviu, Promis Diagnostics, Valar labs, Uroessentials, NRx Pharmaceuticals, Vesica health, Janssen, Immunity Bio, Trigone pharma, Relmada.

## DATA AVAILABILITY

All data are available in the main text or the supplementary materials, code is available on GitHub, sequencing data will be made available on the SRA.

## SUPPLEMENTARY DATA

**Fig. S1.**
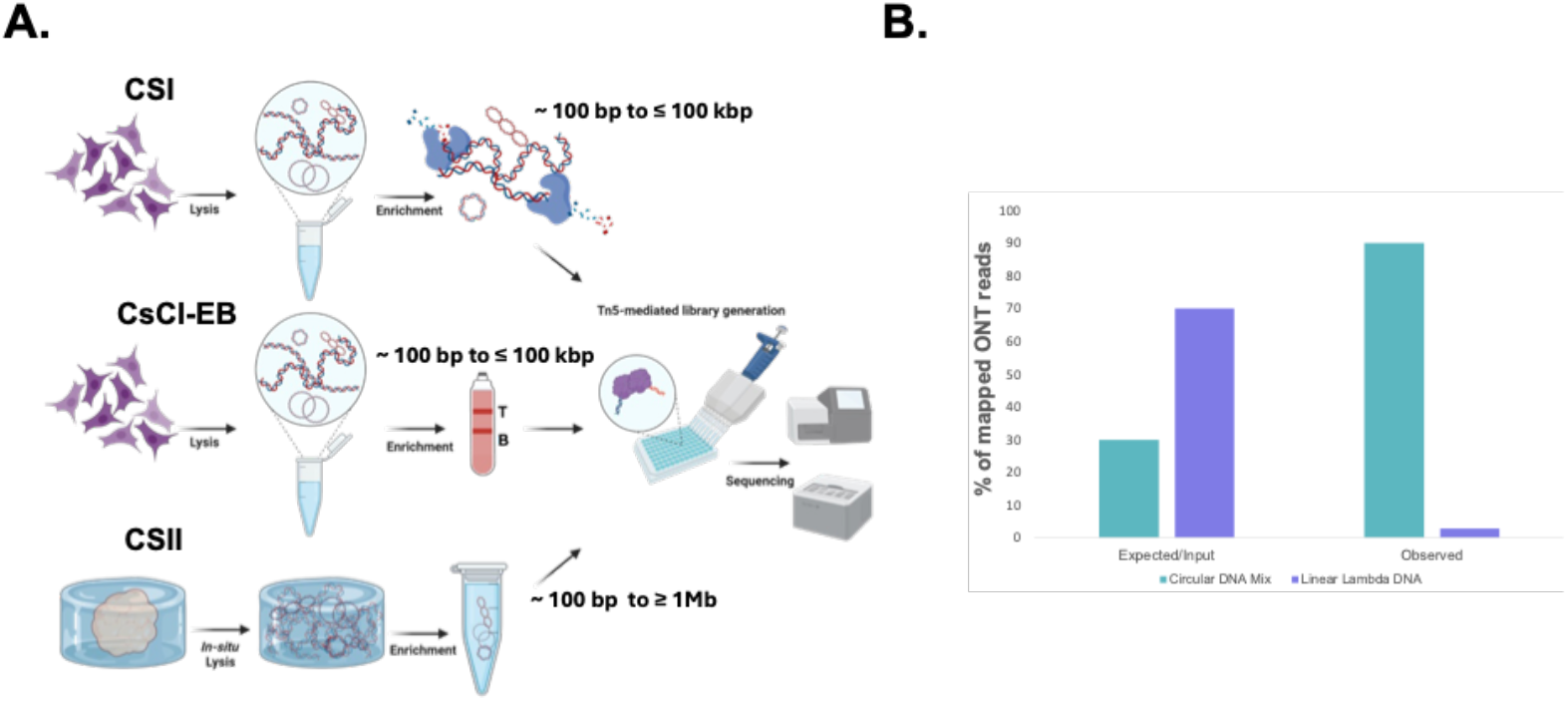
eccDNA isolation/enrichment, sequencing, and analysis workflow. **A.** Circulome-Seq I (CSI) is an enzyme-based workflow for depleting linear DNA, applied directly to genomic DNA without prior processing or selection. CsCl–ethidium bromide (CsCl-EtdBr) density gradient ultracentrifugation separates DNA based on topology: linear DNA and nicked circles band toward the top of the gradient, while covalently closed circular DNAs band toward the bottom. Retrieved DNA from either band can be sequenced without prior amplification. Both CSI and CsCl-EtdBr have inherent size limitations. A third protocol, Circulome-Seq II (CSII) is diagrammed, starting with the embedding of tissues, cells, or other biological samples in a hydrogel matrix containing nucleic acid stabilizers. Genomic DNA can also serve as input for CSII. Sequencing libraries were prepared using Tn5. **B.** Upon CSII enrichment of circular DNA from a mixture of linear lambda DNA and a pool of eccDNA mimics, >95% of the linear lambda DNA is removed (resulting in >95% eccDNA purity); in this analysis, the eccDNA mimics were enriched by >70-fold

**Fig. S2.**
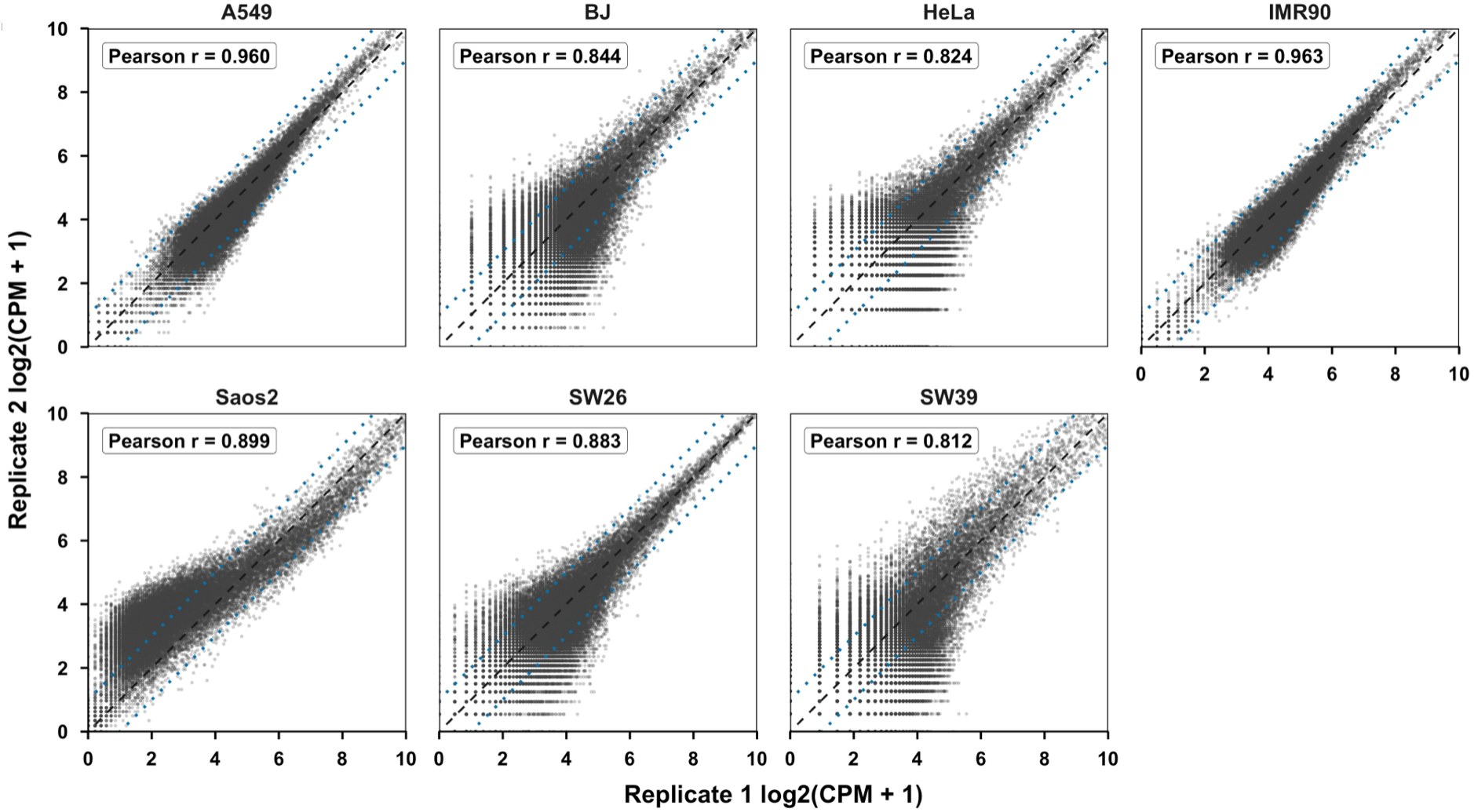
Reproducibility of whole-genome eccDNA read-coverage profiles across replicate preparations. Scatter plots show bin-by-bin comparisons of CPM-normalized eccDNA signal between replicate preparations for each cell line. Each point represents one genomic bin (bin size is 100-kbp.) The x and y axes show log2(CPM + 1) values from replicates 1 and 2, respectively, where CPM denotes counts per million mapped reads and normalizes bin counts by total library size. The dashed diagonal line indicates equal signal between replicates, and the dotted lines indicate two-fold differences. Pearson correlation coefficients, calculated from log2(CPM + 1) values, are shown in each panel.

**Fig. S3.**
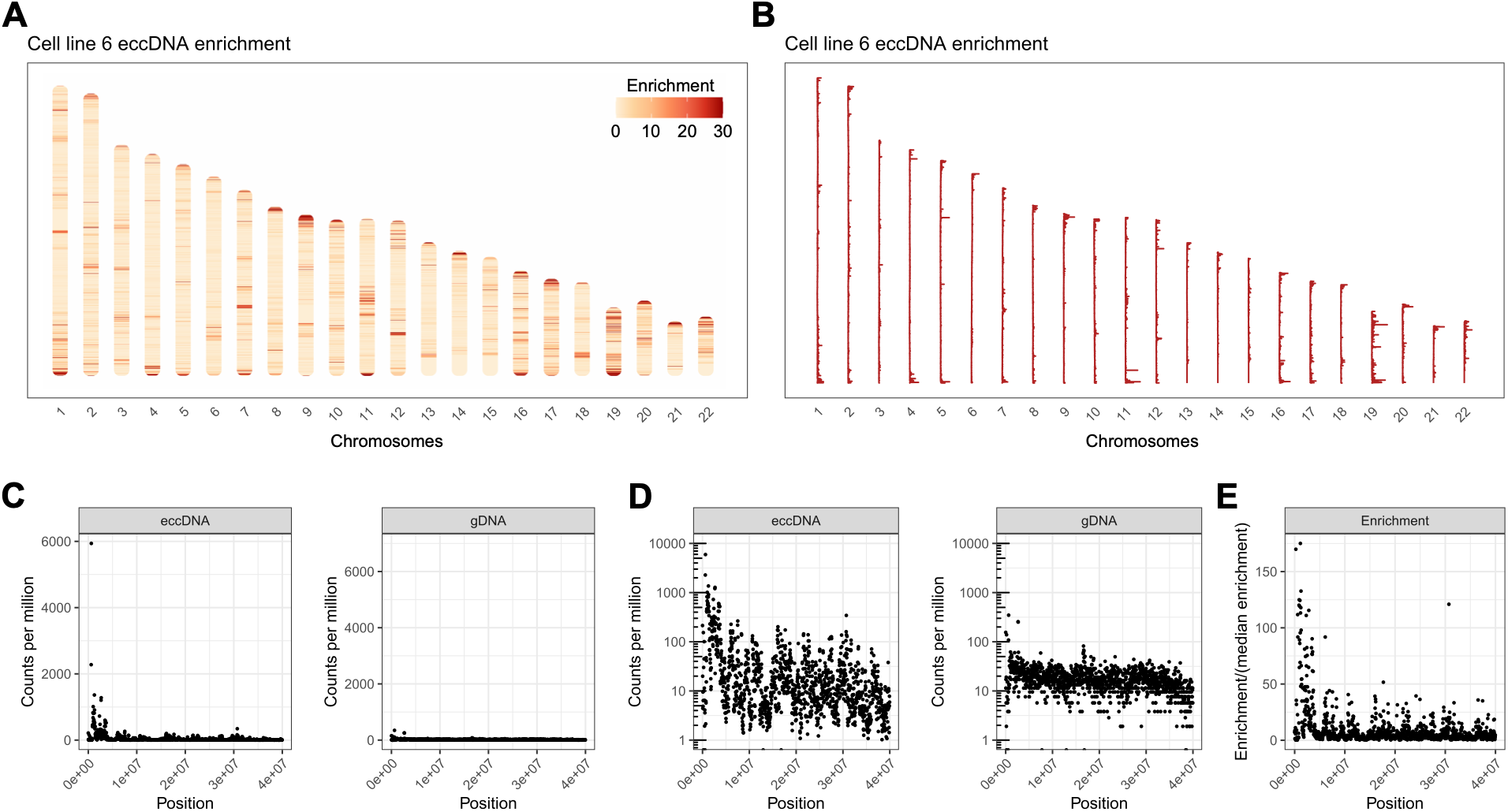
Quantification of eccDNA using short-read alignment with Burrow-Wheeler Aligner (BWA). **(A)** Ideogram of human chromosomes with color scale showing eccDNA enrichment in 30 kb bins. Enrichment at each bin i is estimated as r_i = (eccDNA_i/ gDNA_i) / median_j(eccDNA_j/ gDNA_j). Colors are capped at 30. **(B)** Ideogram of the same data as in (A), plotted as a genome-wide line trace of r_i across chromosomes. **(C)** Scatter plots of eccDNA (left) and gDNA (right) read counts per million (CPM) across the first 40 Mb of chromosome 1, plotted in 30-kb non-overlapping bins. **(D)** Same as (C) with the y-axis on a log10 scale. **(E)** Genome-wide eccDNA enrichment (r_i) as in (A), plotted as a scatter plot of r_i values.

**Fig. S4.**
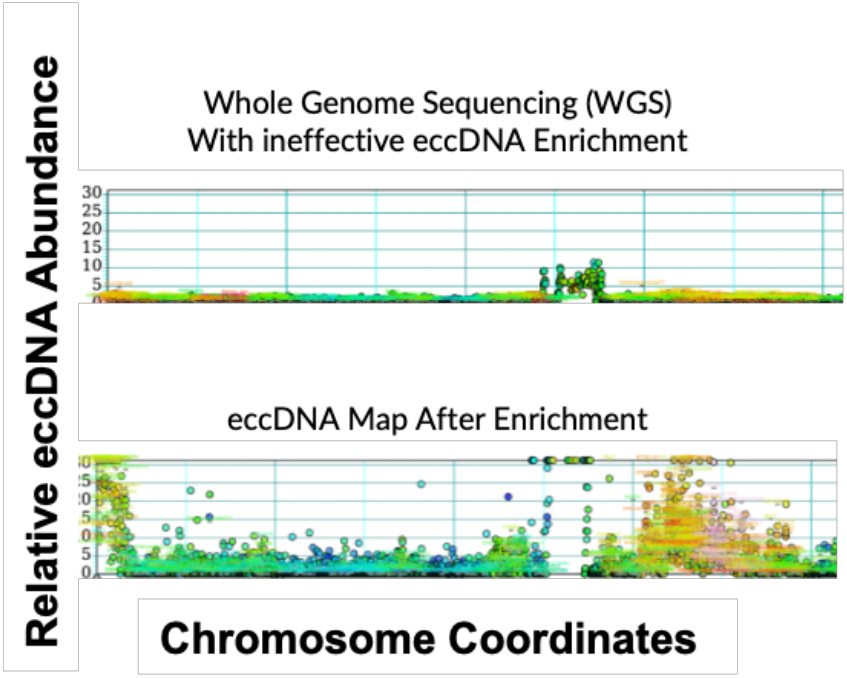
Whole-genome sequencing without effective circular DNA enrichment fails to detect eccDNA due to overwhelming linear DNA background. Top: Normalized eccDNA signal from a controlled unsuccessful enrichment experiment, shown as the ratio of eccDNA reads to matched genomic DNA reads. **Bottom:** The same analysis performed after successful circular DNA enrichment, revealing a clear and robust eccDNA signal. Note that these eccDNA signals were hidden in the top panel due to the high abundance of linear DNA relative to eccDNA in genomic DNA.

